# Role of spike in the pathogenic and antigenic behavior of SARS-CoV-2 BA.1 Omicron

**DOI:** 10.1101/2022.10.13.512134

**Authors:** Da-Yuan Chen, Devin Kenney, Chue Vin Chin, Alexander H. Tavares, Nazimuddin Khan, Hasahn L. Conway, GuanQun Liu, Manish C. Choudhary, Hans P. Gertje, Aoife K. O’Connell, Darrell N. Kotton, Alexandra Herrmann, Armin Ensser, John H. Connor, Markus Bosmann, Jonathan Z. Li, Michaela U. Gack, Susan C. Baker, Robert N. Kirchdoerfer, Yachana Kataria, Nicholas A. Crossland, Florian Douam, Mohsan Saeed

## Abstract

The recently identified, globally predominant SARS-CoV-2 Omicron variant (BA.1) is highly transmissible, even in fully vaccinated individuals, and causes attenuated disease compared with other major viral variants recognized to date^1–7^. The Omicron spike (S) protein, with an unusually large number of mutations, is considered the major driver of these phenotypes^3,8^. We generated chimeric recombinant SARS-CoV-2 encoding the S gene of Omicron in the backbone of an ancestral SARS-CoV-2 isolate and compared this virus with the naturally circulating Omicron variant. The Omicron S-bearing virus robustly escapes vaccine-induced humoral immunity, mainly due to mutations in the receptor-binding motif (RBM), yet unlike naturally occurring Omicron, efficiently replicates in cell lines and primary-like distal lung cells. In K18-hACE2 mice, while Omicron causes mild, non-fatal infection, the Omicron S-carrying virus inflicts severe disease with a mortality rate of 80%. This indicates that while the vaccine escape of Omicron is defined by mutations in S, major determinants of viral pathogenicity reside outside of S.

---

As of March 2022, the successive waves of the coronavirus disease 2019 (COVID-19) pandemic have been driven by five major SARS-CoV-2 variants, called variants of concern (VOC); Alpha (B.1.1.7), Beta (B.1.351), Gamma (P.1), Delta (B.1.617.2 and AY lineages), and Omicron (BA lineages)^9^. Omicron is the most recently recognized VOC that was first documented in South Africa, Botswana, and in a traveler from South Africa in Hong Kong in November 2021 (GISAID ID: EPI_ISL_7605742)^10,11^. It quickly swept through the world, displacing the previously dominant Delta variant within weeks and accounting for the majority of new SARS-CoV-2 infections by January 2022^12–16^. Omicron has at least three lineages, BA.1, BA.2, and BA.3, with the former being the most predominant lineage worldwide^13,17–19^. BA.1 (hereinafter referred to as Omicron) exhibits a remarkable escape from infection- and vaccine-induced humoral immunity^4,5,20,21^. Further, it is less pathogenic than other VOCs in humans and *in vivo* models of infection^1–3,22–26^. Omicron differs from the prototype SARS-CoV-2 isolate, Wuhan-Hu-1, by 59 amino acids; 37 of these changes are in the S protein, raising the possibility that S is at the heart of Omicron’s pathogenic and antigenic behavior.

## Spike mutations only partially affect the replication of Omicron in cell culture

The Omicron S protein carries 30 amino acid substitutions, 6 deletions, and one three-amino acid-long insertion compared to Wuhan-Hu-1 (**Fig. 1a,b**). Twenty-five of these changes are unique to Omicron relative to other VOCs, although some of them have been reported in waste water and minor SARS-CoV-2 variants^27–29^. To test the role of the S protein in Omicron phenotype, we generated a chimeric recombinant virus containing the S gene of Omicron (USA-lh01/2021) in the backbone of an ancestral SARS-CoV-2 isolate (GISAID EPI_ISL_2732373)^30^ (**Fig. 1c**). To produce this chimeric Omi-S virus, we employed a modified form of cyclic polymerase extension reaction (CPER) (**Extended Data Fig. 1**) that yielded highly concentrated virus stocks, containing 0.5-5 × 10^6^ plaque-forming units (PFU) per ml, from transfected cells within two days of transfection (**Fig. 1d,e**), obviating the need for additional viral amplification^31,32^.

**Fig. 1:**
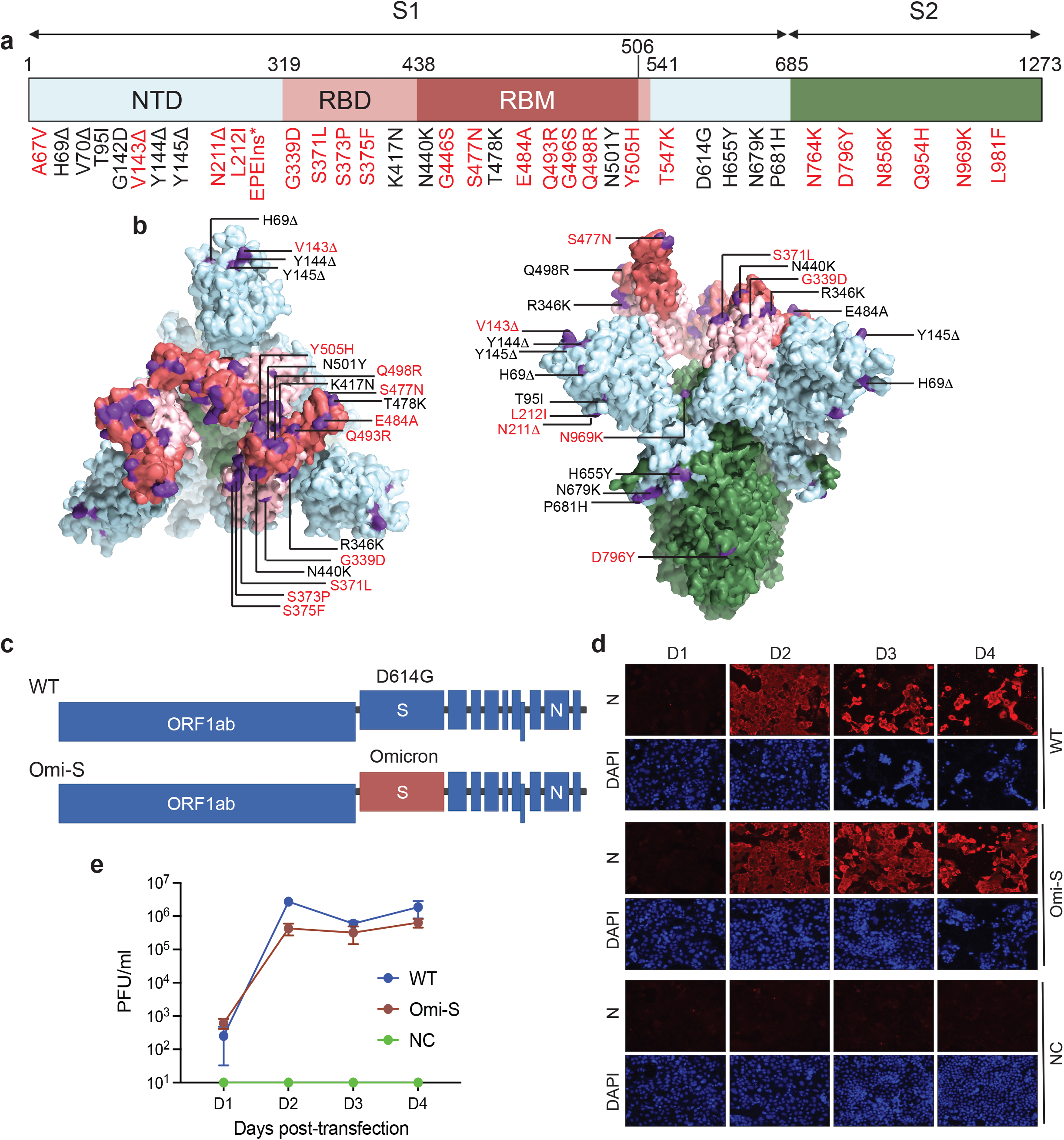
Generating Recombinant SARS-Co-2 by CPER. **a,** Schematic overview of mutations in Omicron spike (in comparison to the SARS-CoV-2 Wuhan-Hu-1 isolate; NCBI accession number: NC_045512). Numbering is based on Wuhan-Hu-1 sequence. Mutations not reported in previous variants of concern are shown in red. NTD, N-terminal domain; RBD, receptor-binding domain; RBM, receptor-binding motif. **b,** Location of Omicron mutations on the trimeric spike protein. Domains are colored according to a. **c,** Schematic of recombinant SARS-CoV-2 generated by CPER. S, spike; N, nucleocapsid. **d,** ACE2/TMPRSS2/Caco-2 cells transfected with the SARS-CoV-2 CPER product were stained with an anti-nucleocapsid antibody on indicated days post-transfection. DAPI was used to stain the cell nuclei. NC, negative control generated by omitting Fragment 9 from the CPER reaction. **e,** Virus titer in the culture medium of the transfected cells at indicated days post-transfection, as measured by the plaque assay. The data are plotted as mean ± SEM of two experimental repeats.

We first compared the infection efficiency of Omi-S with an ancestral virus and Omicron in cell culture (**Fig. 2a**). For this, we infected ACE2/TMPRSS2/Caco-2^33^ and Vero E6 cells with Omi-S, a recombinant D614G-bearing ancestral virus (GISAID EPI_ISL_2732373)^30^, and a clinical Omicron isolate (USA-lh01/2021) at a multiplicity of infection (MOI) of 0.01 and monitored viral propagation by flow cytometry and the plaque assay. The ancestral virus [hereinafter referred to as wild-type (WT)] and Omi-S spread fast in ACE2/TMPRSS2/Caco-2 cells, yielding 89% and 80% infected cells, respectively, at 24 hours post-infection (hpi) (**Fig. 2b**). In contrast, Omicron replicated slower, leading to 48% infected cells at 24 hpi. A similar pattern was seen in Vero E6 cells, where 60% and 41% of cells were positive for WT and Omi-S, respectively, at 48 hpi, in contrast to 10% positive cells for Omicron (**Fig. 2c**). The plaque assay showed that although both Omi-S and Omicron produced lower levels of infectious virus particles compared with WT, the viral titer of Omi-S was significantly higher than that of Omicron. In ACE2/TMPRSS2/Caco-2 cells, Omi-S produced 5.1-fold (*p* = 0.0006) and 5.5-fold (*p* = 0.0312) more infectious particles than Omicron at 12 hpi and 24 hpi, respectively (**Fig. 2d**). Similarly, in Vero E6 cells, the infectious virus titers of Omi-S were 17-fold (*p* = 0.0080) and 11-fold (*p* = 0.0078) higher than that of Omicron at 24 hpi and 48 hpi, respectively (**Fig. 2e**). The difference between viruses became less obvious at later time points due to higher cytotoxicity caused by Omi-S compared with Omicron (**Fig. 2f**). The higher infection efficiency of Omi-S relative to Omicron was also reflected in the plaque size; while WT produced the largest plaques (~ 4.1 mm), the size of Omi-S plaques (~2.2 mm) was 2-fold (*p* < 0.0001) larger than that of Omicron plaques (~1.1 mm) (**Fig. 2g**). These results indicate that while mutations in the S protein influence the infection efficiency of Omicron, they do not fully explain the infection behavior of Omicron in cell culture.

**Fig. 2:**
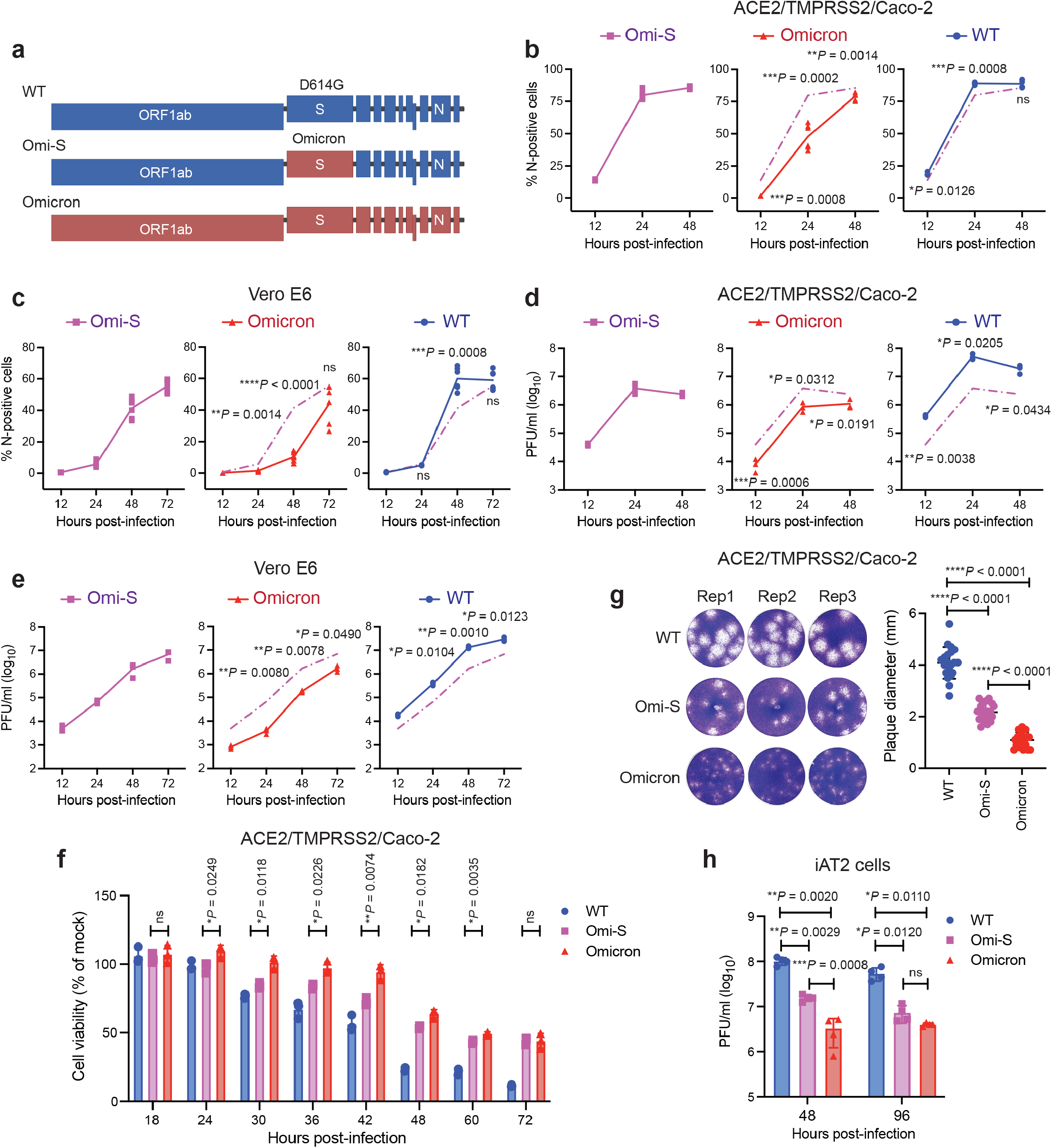
Effect of spike on *in vitro* growth kinetics of Omicron. **a**, Schematic of viruses used in this figure. S, spike; N, nucleocapsid. **b-e,** ACE2/TMPRSS2/Caco-2 and Vero E6 cells were infected at an MOI of 0.01, and the percentage of nucleocapsid (N)-positive cells (n = 6) (**b,c**) and levels of infectious virus production (n = 3) (**d,e**) were determined by flow cytometry and the plaque assay, respectively. **f,** The cell viability of SARS-CoV-2-infected ACE2/TMPRSS2/Caco-2 cells (MOI of 0.1) was quantified by the CellTiter-Glo assay at indicated time points. The *P* values reflect a statistically significant difference between Omi-S and Omicron. **g,** Plaque sizes. Left, representative images of plaques on ACE2/TMPRSS2/Caco-2 cells. Right, diameter of plaques is plotted as mean ± SD of 20 plaques per virus. **h,** Human induced pluripotent stem cell-derived alveolar type 2 epithelial cells were infected at an MOI of 2.5 for 48h or 96h. The apical side of cells was washed with 1X PBS and the levels of infectious virus particle were measured by the plaque assay. n = 4. Data are mean ± SD from the indicated number of biological replicates. Experiments were repeated twice, with each experimental repeat containing 2 (**h**) or 3 (**b-g**) replicates. *p* values were calculated by a two-tailed, unpaired *t*-test with Welch’s correction. \**p* <0.05, \*\**p* <0.01, \*\*\**p* <0.001, and \*\*\*\**p* < 0.0001; ns, not significant.

We next expanded our studies to lung epithelial cells, which are a major viral replication site in patients with severe COVID-19. Accordingly, we employed human induced pluripotent stem cell-derived lung alveolar type 2 epithelial (iAT2) cells. AT2 cells represent an essential cell population in the distal lung and constitute one of the primary targets of SARS-CoV-2 infection^34–36^. We infected iAT2 cells, grown as an air-liquid interface (ALI) culture, at an MOI of 2.5 and monitored the secretion of viral progeny on the apical interface of cells at 48 hpi and 96 hpi. In congruence with the results obtained from cell lines, WT SARS-CoV-2 produced the highest levels of infectious virus particles (**Fig. 2h**). Among the Omi-S and Omicron, the former yielded ~5-fold (*p* = 0.0008) higher infectious viral titer at 48 hpi. The viral titers for WT and Omi-S decreased at 96 hpi compared with 48 hpi due to the cytopathic effect (CPE) of infection. However, no CPE was seen for Omicron, leading to sustained production of infectious virions. Overall, these results corroborate the conclusion that mutations in S do not fully account for the attenuated replication capacity of Omicron in cultured human cells.

## Spike has an appreciable but minimal role in Omicron pathogenicity in K18-hACE2 mice

To examine if Omi-S exhibits higher *in vivo* fitness compared with Omicron, we investigated the infection outcome of Omi-S relative to WT SARS-CoV-2 and Omicron in K18-hACE2 mice. In agreement with the published literature^3,37–39^, intranasal inoculation of mice (aged 12-20 weeks) with Omicron (10^4^ PFU per animal) caused no significant weight loss, whereas inoculation with WT virus triggered a rapid decrease in body weight with all animals losing over 20% of their initial body weight by 8 days post-infection (dpi) (**Fig. 3a**). Importantly, 80% of animals infected with Omi-S also lost over 20% of their body weight by 9 dpi (**Fig. 3a and Extended Data Fig. 2a**). The evaluation of clinical scores (a cumulative measure of weight loss, abnormal respiration, aberrant appearance, reduced responsiveness, and altered behavior) also revealed a similar pattern; while Omicron-infected mice displayed little to no signs of clinical illness, the health of those infected with WT and Omi-S rapidly deteriorated, with the former inflicting a more severe disease (*p* = 0.0102) (**Fig. 3b and Extended Data Fig. 2b**). Since SARS-CoV-2 causes fatal infection in K18-hACE2 mice^3,40,41^, we leveraged this situation to compare the animal survival after viral infection. In agreement with the results of body-weight loss and clinical score, WT and Omi-S caused mortality rates of 100% (6/6) and 80% (8/10), respectively. In contrast, all animals infected with Omicron survived (**Fig. 3c**). These findings indicate that the S protein is not the primary determinant of Omicron’s pathogenicity in K18-hACE2 mice.

**Fig. 3:**
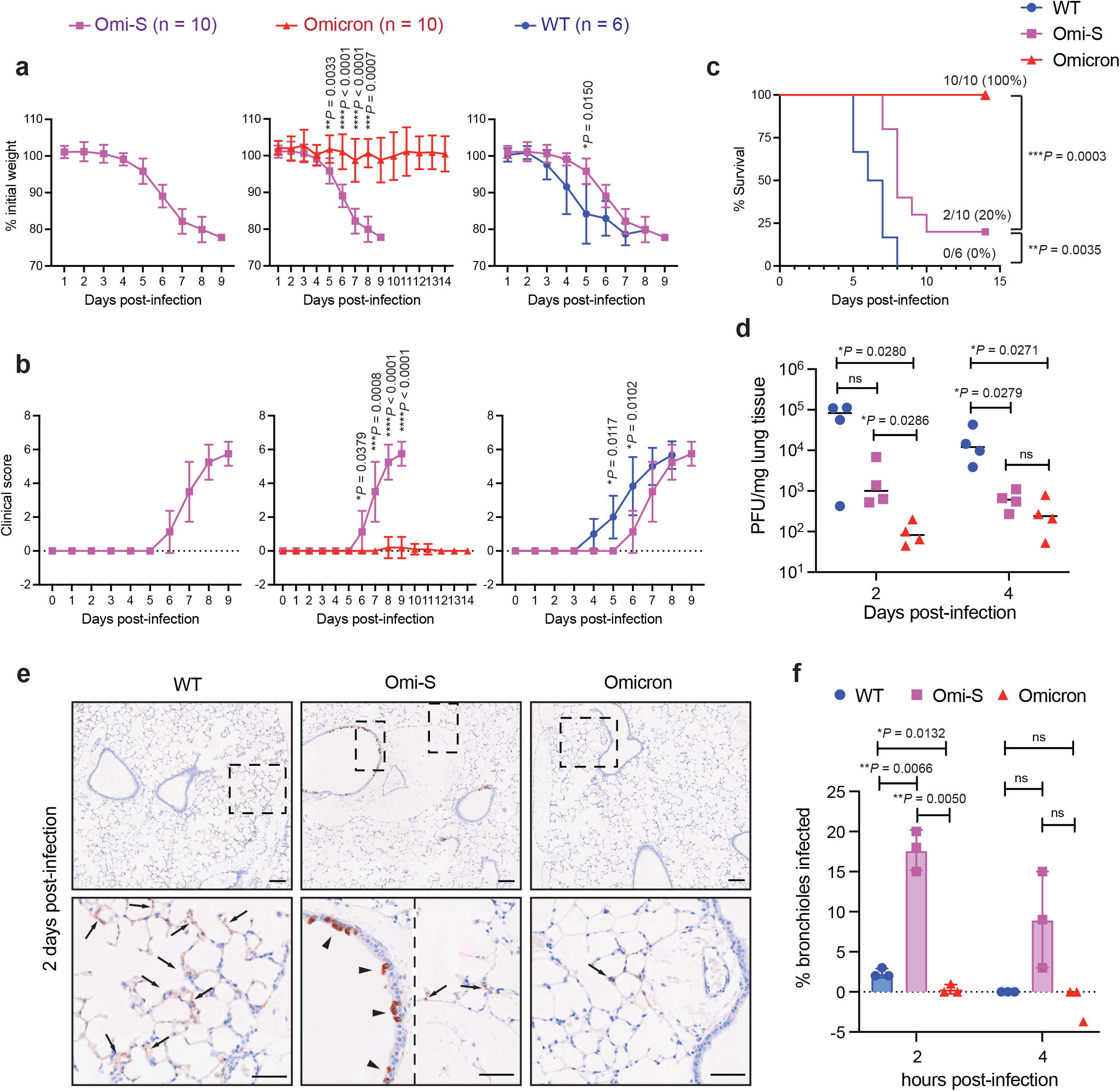
Role of spike in Omicron pathogenicity. **a-c,** Male and female K18-hACE2 mice (aged 12-20 weeks) were intranasally inoculated with 1 × 10^4^ PFU of WT (n = 6), Omi-S (n = 10), or Omicron (n = 10). Two independently generated virus stocks were used in this experiment. The body weight (**a**), clinical score (**b**), and survival (**c**) were monitored daily for 14 days. Animals losing 20% of their initial body weight were euthanized. **d,e,** K18-hACE2 mice were intranasally inoculated with 1 × 10^4^ PFU of WT (n = 7), Omi-S (n = 7), and Omicron (n = 7). Lung samples of the infected mice were collected at 2 or 4 dpi to determine the viral titer (n = 4) (**d**) or for immunohistochemistry (IHC) detection of the S protein (n = 3) (**e**). In e, representative images of IHC staining for the detection of the SARS-CoV-2 S protein (brown color) in alveoli (arrows) and bronchioles (arrowheads) in the lungs of the infected mice at 2 dpi are shown. (Scale bar = 100 μm). **f,** The percentage of S-positive bronchioles in the lungs of infected mice. Each dot represents an infected animal. Data are presented as mean ± SD from the indicated number of biological replicates. Statistical significance was determined using two-tailed, unpaired *t*-test with Welch’s correction (**a,b,d,f**) and log-rank (Mantel-Cox) test (**c**). \**p* <0.05, \*\**p* <0.01, \*\*\**p* <0.001, and \*\*\*\**p* < 0.0001; ns, not significant.

Next, we compared the virus propagation of Omi-S with Omicron and WT SARS-CoV-2 in the lungs of K18-hACE2 mice. The mice (12-20 weeks old) were intranasally challenged with 10^4^ PFU (7 mice per virus), and their lungs were collected at 2 and 4 dpi for virological and histological analysis. Consistent with *in vitro* findings, the infectious virus titer in the lungs of WT-infected mice was higher than that detected in mice infected with other two viruses (**Fig. 3d**). Notably however, Omi-S-infected mice produced 30-fold (*p* = 0.0286) more infectious virus particles compared with Omicron-infected mice at 2 dpi. The titer decreased at 4 dpi for WT- and Omi-S-infected mice, yet it showed an increasing trend for Omicron-infected animals, pointing to the possibility of mild but persistent infection by Omicron in K18-hACE2 mice.

To evaluate the viral pathogenicity in the lungs, we performed histopathological analysis of the lung tissue of infected K18-hACE2 mice. As previously reported^3,42^, an extensive near-diffused immunoreactivity of the SARS-CoV-2 S protein was detected in lung alveoli of mice infected with WT virus (**Fig. 3e**). In contrast, Omi-S and Omicron infection produced localized foci of alveolar staining with fewer foci for Omicron compared with Omi-S. The most striking phenotype was seen in bronchiolar epithelium. While Omi-S virus caused a severe bronchiolar infection with around 15-20% of bronchioles being positive for the S protein in all mice examined at 2 dpi, less than 1% bronchioles were S-positive in Omicron-infected mice (**Fig. 3f**). Further, bronchiolar infection was associated with epithelial necrosis in Omi-S-infected mice, as determined through serial hematoxylin and eosin (H&E) section analysis, whereas no histological evidence of airway injury was observed in Omicron-infected mice (**Extended Data Fig. 3**). This suggests that the replication of Omicron in mice lungs, particularly in bronchioles, is substantially attenuated compared with Omi-S, supporting our conclusion that mutations in the S protein are only partially responsible for the attenuated pathogenicity of Omicron.

## Mutations in the spike RBM are major drivers of Omicron’s escape from neutralization

Next, we examined if Omi-S captures the immune escape phenotype of Omicron. A large body of literature has demonstrated extensive escape of Omicron from vaccine-induced humoral immunity ^4,10,43^. We compared the *in vitro* neutralization activity of sera obtained from vaccinated individuals against the SARS-CoV-2 Washington isolate (USA-WA1/2020), Omi-S, and Omicron. Sera collected within two months of the second dose of mRNA-1273 (Moderna mRNA vaccine; n = 12) or BNT162b2 (Pfizer-BioNTech mRNA vaccine; n = 12) vaccine were included (**Extended Data Table 1**). We performed a multicycle neutralization assay using a setting in which the virus and neutralizing sera were present at all times, mimicking the situation in a seropositive individual. All sera poorly neutralized Omicron, with 11.1-fold (range: 4.4- to 81.2-fold; *p* < 0.0001) lower half-maximal neutralizing dilution (ND_50_) for Omicron compared with WA1 (**Fig. 4a,b**). In fact, around 80% of samples failed to completely neutralize Omicron at the highest tested concentration (**Extended Data Fig. 4**). Notably, Omi-S exhibited identical ND50 values to Omicron (11.5-fold lower than that of WA1; *p* < 0.0001) (**Fig. 4a,b**), suggesting that the Omicron S protein, when incorporated into a WT virus, behaves the same way as in Omicron.

**Fig. 4:**
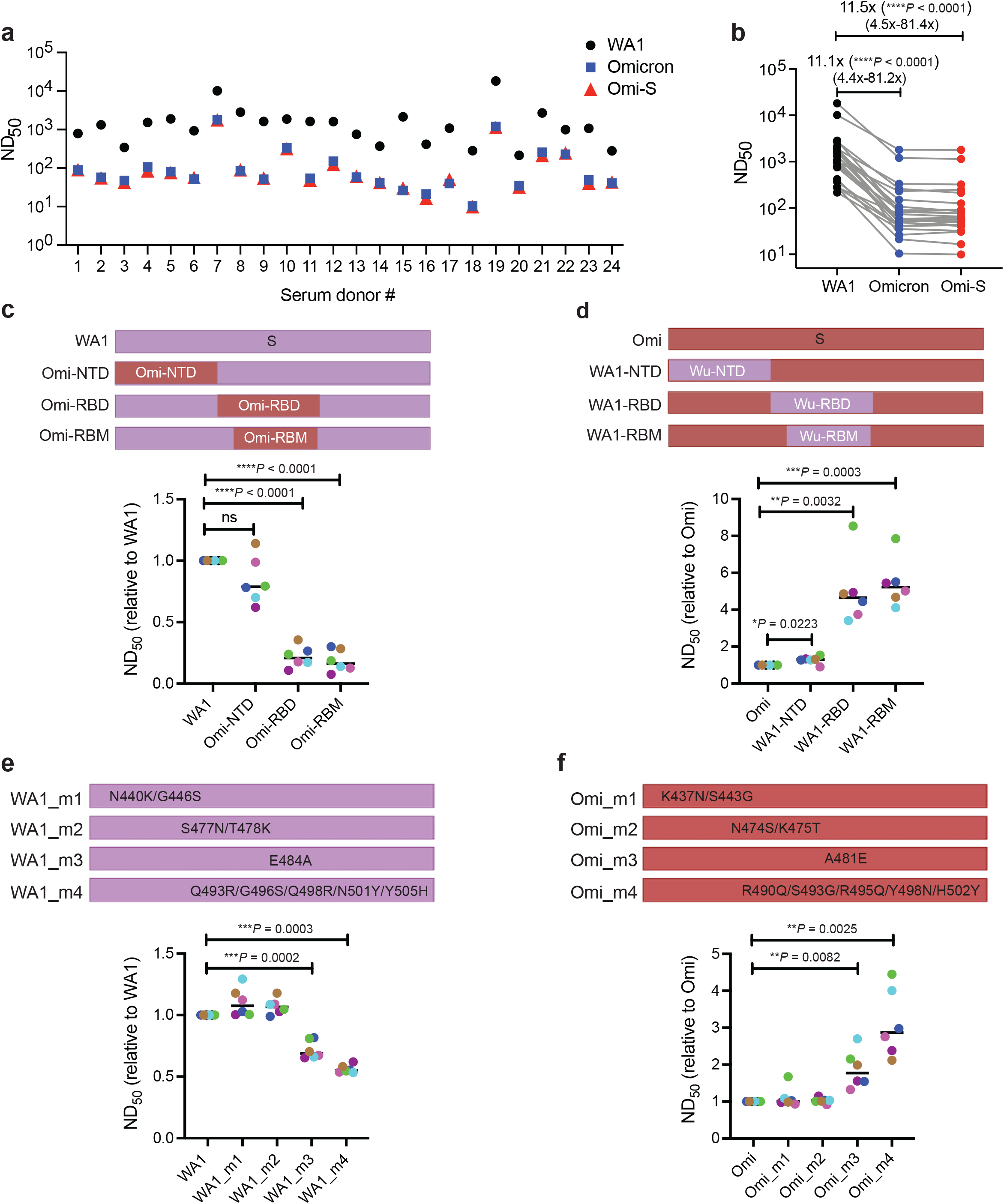
Role of spike in immune resistance of Omicron. **a,** ND_50_ values for WA1, Omi-S, and Omicron in sera from individuals who received two shots of Moderna (donor 1-12) or Pfizer (donor 13-24) vaccine (further details of sera are provided in Extended Data Table 1; individual curves are shown in Extended Data Fig. 4). **b,** Trajectories of ND_50_ values against WA1, Omi-S, and Omicron (the data from a is plotted). Fold-change in ND_50_ values is indicated. **c,d,e,f,** Schematic of the chimeric (**top panels; c,d**) and mutant (**top panels; e,f**) viruses. The amino acid numbering for WA1 mutants in e is based on the WA1 spike sequence, whereas the numbering for Omicron mutants in f is based on the Omicron spike sequence. Six of the 24 sera (three from Moderna and three from Pfizer) were tested. Each serum sample is represented by a dot of specific color. The data are plotted as fold-change of the parental virus. Statistical significance was determined using a two-tailed, unpaired *t* test with Welch’s correction. \**p* <0.05, \*\**p* <0.01, \*\*\**p* <0.001, and \*\*\*\**p* < 0.0001; ns, not significant.

The SARS-CoV-2 S protein comprises two domains: the S1 domain, which interacts with the ACE2 receptor, and the S2 domain, which is responsible for membrane fusion^44^. Within the S1 domain lie an N-terminal domain (NTD) and a receptor-binding domain (RBD), which harbors the receptor-binding motif (RBM) that makes direct contact with the ACE2 receptor^45^. The NTD of Omicron S carries 11 amino acid changes, including 6 deletions and one three-amino acid-long insertion, whereas the RBD harbors 15 mutations, 10 of which are concentrated in the RBM (**Fig. 1a,b**). Both NTD and RBD host neutralizing epitopes^46–50^, but the RBD is immunodominant and represents the primary target of the neutralizing activity present in SARS-CoV-2 immune sera^50,51^. To determine if the neutralization resistance phenotype of Omicron is caused by mutations in a particular S domain, we generated two groups of chimeric viruses. The first group comprised the WA1 virus carrying the NTD, RBD, or RBM of Omicron (**Fig. 4c**), and the second group consisted of Omi-S virus bearing the NTD, RBD, or RBM of WA1 (**Fig. 4d**). The neutralization assay showed that mutations in the RBM were the major cause of Omicron’s resistance to vaccine-induced humoral immunity: replacing the RBM of WA1 with that of Omicron decreased ND_50_ by 5.4-fold (*p* < 0.0001), and conversely, substituting the RBM of Omi-S with that of WA1 increased ND_50_ by 5.6-fold (*p* = 0.0003) (**Fig. 4c,d**). The fact that none of the RBM-swap viruses achieved the difference of ~11-fold seen between WA1 and Omi-S suggests that mutations in other parts of S also contribute to vaccine resistance.

To investigate if specific mutations in Omicron RBM drive vaccine escape, we generated two additional panels of recombinant viruses, one with WA1 spike carrying Omicron RBM mutations, either singly or in combination (**Fig. 4e**), and the other with Omicron spike lacking the same set of mutations (**Fig. 4f**). Two WA1 mutants, mutant 3 (carrying E484A substitution) and mutant 4 (bearing a cluster of five substitutions Q493R, G496S, Q498R, N501Y, Y505H) exhibited a moderate but statistically significant decrease of 1.4-fold (*p* = 0.0002) and 1.8-fold (*p* = 0.0003) in ND50 values, respectively, compared with WA1 (**Fig. 4e**). The opposite was observed when these mutations were removed from Omicron S; the Omicron mutant 3 (lacking E484A substitution) and mutant 4 (lacking Q493R, G496S, Q498R, N501Y, Y505H) had a 1.9-fold (p = 0.0082) and 3.1-fold (p = 0.0025) higher ND_50_ values compared with Omicron (**Fig. 4f**). Since none of the mutants captured the overall phenotype of Omicron, we assume that the vaccine escape is a cumulative effect of mutations distributed along the length of the S protein. It is possible that mutations alter the conformation of Omicron S in such a manner that most of the immunodominant neutralizing epitopes are disrupted and become unavailable for neutralization.

## DISCUSSION

This study provides important insights into Omicron pathogenicity. We show that spike, the single most mutated protein in Omicron, has an incomplete role in Omicron attenuation. In *in vitro* infection assays, the Omicron spike-bearing ancestral SARS-CoV-2 (Omi-S) exhibits much higher replication efficiency compared with Omicron. Similarly, in K18-hACE2 mice, Omi-S contrasts with non-fatal Omicron and causes a severe disease leading to around 80% mortality. This suggests that mutations outside of spike are major determinants of the attenuated pathogenicity of Omicron in K18-hACE2 mice. Further studies are needed to identify those mutations and decipher their mechanisms of action.

One potential limitation of our study is the use of K18-hACE2 mice for pathogenesis studies instead of the primate models that have more similarities with humans^52,53^. It should however be noted that the K18-hACE2 mouse model is a well-established model for investigating the lethal phenotype of SARS-CoV-2^3,42,54–56^. While these mice develop lung pathology following SARS-CoV-2 infection, mortality has been associated with central nervous system involvement due to viral neuroinvasion^42,57^. The fact that infection with Omi-S, but not with Omicron, elicits neurologic signs, such as hunched posture and lack of responsiveness, in K18-hACE2 mice suggests that the neuroinvasion property is preserved in Omi-S, and the determinants of this property lie outside of the spike protein.

We found that while the ancestral virus mainly replicates in lung alveoli and causes only rare infection of bronchioles in K18-hACE2 mice, Omi-S with isogenic ancestral virus backbone exhibits higher propensity to replicate in bronchiolar epithelium. This is consistent with a hamster study demonstrating higher predilection of Omicron for bronchioles^1^. In vitro studies have also showed that while Omicron replicates poorly in lower lung cells^58^, it causes a robust infection in bronchiolar and nasal epithelial cells^58–60^. Our findings indicate that the higher preference of Omicron for bronchioles is dictated by mutations in the spike protein. We speculate that both Omi-S and Omicron enter the bronchiolar epithelium of K18-hACE2 mice, yet only Omi-S replicates to high enough levels to manifest in overt bronchiolar injury. The preference of Omicron spike for bronchiolar epithelium is likely mediated by its improved efficiency to utilize Cathepsin B/L^58–62^, which form an active viral entry pathway in bronchioles and other airway cells^59,63^. In contrast, SARS-CoV-2 entry into alveolar epithelial cells is mainly driven by TMPRSS2^36,64^, which Omicron spike is deficient in utilizing^60,65^, leading to poor infection of these cells^3,37,58,60^. These findings explain the higher transmission and lower lung pathology caused by Omicron.

Our study shows that mutations in the RBM of Omicron spike are the major determinants of the viral escape from neutralizing antibodies, although mutations in other regions of spike also contribute. Within the RBM, we identify two hotspots of mutations, which impart on Omicron spike the ability to resist neutralization: one bearing the E484A substitution and the other harboring a cluster of five substitutions, Q493R, G496S, Q498R, N501Y and Y505H. The E484A substitution has been shown to escape neutralization by convalescent sera^66^. Further, structural modeling suggests that some therapeutic monoclonal antibodies establish highly stable salt bridges with the E484 residue, entirely losing their binding when this residue is changed to A or upon Q493K and Y505H changes^67^. Similarly, mapping of RBM residues that directly interact with 49 known neutralizing antibodies revealed N440, G446, S477, and T478 as low-frequently interactors, N501, Y505, and Q498 as medium-frequency interactors, and E484 and Q493 as high-frequency interactors^68^, which is in line with our neutralization assay results. Interestingly, while antibody-binding potential of Omicron spike is impaired^69^, its receptor-binding capacity is intact. In fact, the Omicron RBD has higher affinity for ACE2 relative to the Wuhan-Hu-1 and Delta RBDs^60^. This indicates that mutations in the Omicron spike have evolved in such a manner that they hinder antibody binding but preserve the receptor engagement. This opens up the possibility of targeting the conserved and structurally constrained regions of spike involved in ACE2 recognition for the design of broad-spectrum vaccines to control the current COVID-19 pandemic.

## MATERIALS AND METHODS

### Cells, antibodies, and plasmids

The cell lines were incubated at 37°C and 5% CO_2_ in a humidified incubator. Human embryonic kidney HEK293T cells (ATCC; CRL-3216), human lung adenocarcinoma A549 cells (ATCC; CCL-185), human colorectal adenocarcinoma Caco-2 cells (ATCC; HTB-37), and African green monkey kidney Vero E6 cells were maintained in DMEM (Gibco; #11995-065) containing 10% FBS and 1X non-essential amino acids. Lentiviral delivery system was used to generate cells stably expressing human ACE2 and TMPRSS2. Mycoplasma negative status of all cell lines was confirmed.

Anti-SARS-CoV nucleocapsid (N) protein antibody (Rockland; #200-401-A50) was used for detection of the SARS-CoV-2 N protein by IF. Expression plasmid encoding the spike protein of the SARS-CoV-2 Wuhan isolate, pCSII-SARS-CoV-2 F8, was a kind gift from Yoshiharu Matsuura^32^. We replaced the Wuhan spike in this plasmid with a chemically synthesized version of Omicron spike and called the resulting plasmid pCSII-SARS-CoV-2 F8_Omicron. The lentiviral vectors, pLOC_hACE2_PuroR and pLOC_hTMPRSS2_BlastR, containing human ACE2 and TMPRSS2, respectively, have been previously described^33^.

### Omicron stock preparation and titration

All procedures were performed in a biosafety level 3 (BSL3) facility at the National Emerging Infectious Diseases Laboratories of the Boston University using biosafety protocols approved by the institutional biosafety committee (IBC). The SARS-CoV-2 BA.1 Omicron virus stock was generated in ACE2/TMPRSS2/Caco-2 cells. Briefly, 5 × 10^5^ cells, grown overnight in DMEM/10%FBS/1X NEAA in one well of a 6-well plate, were inoculated with the collection medium in which the nasal swab from a SARS-CoV-2 patient was immersed. The swab material was obtained from the Department of Public Health, Massachusetts, and it contained the sequence-verified Omicron virus (NCBI accession number: OL719310). Twenty-four hours after infecting cells, the culture medium was replaced with 2 ml of DMEM/2%FBS/1X NEAA and the cells were incubated for another 72h, at which point the CPE became visible. The culture medium was harvested, passed through a 0.45 μ filter, and kept at −80°C as a P0 virus stock. To generate a P1 stock, we infected 1 × 10^7^ ACE2/TMPRSS2/Caco-2 cells, seeded the day before in a T175 flask, with the P0 virus at an MOI of 0.01. The next day, the culture medium was changed to 25 ml of 2% FBS-containing medium. Three days later, when the cells exhibited excessive CPE, the culture medium was harvested, passed through a 0.45 μ filter, and stored at −80°C as a P1 stock.

To titrate the virus stock, we seeded ACE2/TMPRSS2/Caco-2 cells into a 12-well plate at a density of 2 × 10^5^ cells per well. The next day, the cells were incubated with serial 10-fold dilutions of the virus stock (250 μl volume per well) for 1h at 37°C, overlayed with 1 ml per well of medium containing 1:1 mixture of 2X DMEM/4% FBS and 1.2% Avicel (DuPont; RC-581), and incubated at 37°C for another three days. To visualize the plaques, the cell monolayer was fixed with 4% paraformaldehyde and stained with 0.1% crystal violet, with both fixation and staining performed at room temperature for 30 minutes each. The number of plaques were counted and the virus titer was calculated.

### Recombinant SARS-CoV-2 generation by CPER

SARS-CoV-2 recombinant viruses were generated by using a modified form of the recently published CPER protocol^32,70^. Full-length SARS-CoV-2 cDNA cloned onto a bacterial artificial chromosome (BAC)^30^ was used as a template to amplify the viral genome into eight overlapping fragments (F1, F2, F3, F4, F5, F6, F7, and F9). The pCSII-SARS-CoV-2 F8 and pCSII-SARS-CoV-2 F8_Omicron plasmids, which were used to generate spike mutants, served as templates for amplification of fragment 8 (F8). A UTR linker containing a hepatitis delta virus ribozyme (HDVr), the bovine growth hormone polyadenylation signal sequence (BGH-polyA), and a cytomegalovirus (CMV) promoter was cloned onto a pUC19 vector and used as a template to amplify the linker sequence. The 5’ termini of all ten DNA fragments (F1-F9 and the linker) were phosphorylated by using T4 PNK (NEB; #M0201), and the equimolar amounts (0.05 pmol each) of the resulting fragments were subjected to a CPER reaction in a 50 μl volume using 2 μl of PrimeStar GXL DNA polymerase (Takara Bio; #R050A). The following cycling conditions were used for CPER: an initial denaturation at 98°C for 2 min; 35 cycles of denaturation at 98°C for 10 s, annealing at 55°C for 15 s, and extension at 68°C for 15 min; and a final extension at 68°C for 15 min. The nicks in the circular product were sealed by using DNA ligase.

To transfect cells with the CPER product, we seeded ACE2/TMPRSS2/Caco-2 cells into a 6-well plate at a density of 5 x10^5^ cells per well. The transfection mix was prepared by mixing 26 μl of the original 52 μl CPER reaction volume with 250 μl of Opti-MEM (Thermo Fisher Scientific; #31985070) and 6 μl of TransIT-X2 Dynamic Delivery System (Mirus Bio; #MIR 6000). Following incubation at room temperature for 25 min, the transfection mix was added to the cells. The next day, the culture medium was replaced with fresh DMEM containing 2% FBS. The CPE became visible in 3-4 days, at which point the culture medium was collected and stored as a P0 virus stock. The P0 stock was used for experiments described in this manuscript. The spike region of all CPER-generated viruses was sequenced by either Sanger sequencing or next generation sequencing to confirm the presence of desired and the absence of adventitious changes.

### SARS-CoV-2 neutralization assay

For neutralization assays, initial 1:10 dilutions of plasma, obtained from individuals who received two shots of either Moderna or Pfizer mRNA-based SARS-CoV-2 vaccine, were five-fold serial diluted in Opti-MEM over seven or eight dilutions. These plasma dilutions were then mixed at a 1:1 ratio with 1.25 × 10^4^ infectious units of SARS-CoV-2 and incubated for 1h at 37°C. Thereafter, 100 μl of this mixture was directly applied to ACE2/A549 cells seeded the previous day in poly-L-lysine-coated 96-well plates at a density of 2.5 × 10^4^ cells per well in 100 μl volume. Thus, the final starting dilution of plasma was 1:20 and the final MOI was 0.5. The cells were incubated at 37°C for 24h, after which they were fixed and stained with an anti-nucleocapsid antibody. When PBS instead of plasma was used as a negative control, these infection conditions resulted in around 40-50% infected cells at 24 hpi.

### Generation and infection of iAT2 cells

The detailed protocol for generation of human iPSC-derived alveolar epithelial type II cells (iAT2s) has been published in our recent papers^36,71^. The air-liquid interface (ALI) cultures were established by preparing single cell suspensions of iAT2 3D sphere cultures grown in Matrigel. Briefly, Matrigel droplets containing iAT2 spheres were dissolved in 2 mg/ml dispase (Sigma) and the spheres were dissociated in 0.05% trypsin (GIBCO) to generate a single-cell suspension. 6.5 mm Transwell inserts (Corning) were coated with dilute Matrigel (Corning) in accordance with the manufacturer’s protocol. Single-cell iAT2s were plated on Transwells at a density of 520,000 cells/cm2 in 100 μl of CK+DCI medium containing 10 μM of Rho-associated kinase inhibitor (“Y”; Sigma Y-27632). 600 μl of this medium was added to the basolateral compartment. 24h after plating, the basolateral medium was changed with fresh CK+DCI+Y medium. 48h after plating, the apical medium was aspirated to initiate ALI culture. 72h after plating, basolateral medium was replaced with CK+DCI medium to remove the rho-associated kinase inhibitor. Basolateral medium was changed every two days thereafter. The detailed composition of CK+DCI medium is provided in our previous publications^36,71^.

iAT2 cells in ALI cultures were infected with purified SARS-CoV-2 stock at an MOI of 2.5 based on the titration done on ACE2/TMPRSS2/Caco-2 cells. For infection, 100 μl of inoculum prepared in 1X PBS (or mock-infected with PBS-only) was added to the apical chamber of each Transwell and incubated for 2h at 37°C followed by the removal of the inoculum and washing of the apical side three times with 1X PBS (100 μl/wash). The cells were incubated for two or four days, after which the newly released virus particles on the apical side were collected by adding 100 μl of 1X PBS twice to the apical chamber and incubating at 37°C for 15 min. The number of infectious virus particles in the apical washes were measured by the plaque assay on ACE2/TMPRSS2/Caco-2 cells. For flow cytometry, iAT2 cells were detached by adding 0.2 ml Accutase (Sigma; #A6964) apically and incubated at room temperature for 15 min. The detached cells were pelleted by low-speed centrifugation, fixed in 10% formalin, and stained with anti-SARS-CoV-2 N antibody.

### Mice maintenance and approvals

Mice was maintained in a facility accredited by the Association for the Assessment and Accreditation of Laboratory Animal Care (AAALAC). Animal studies were performed following the recommendations in the Guide for the Care and Use of Laboratory Animals of the National Institutes of Health. The protocols were approved by the Boston University Institutional Animal Care and Use Committee (IACUC). Heterozygous K18-hACE2 C57BL/6J mice (Strain 2B6.Cg-Tg(K18-ACE2)2Prlmn/J) were purchased from the Jackson Laboratory (Jax, Bar Harbor, ME). Animals were housed in ventilated cages (Tecniplast, Buguggiate, Italy) and maintained on a 12:12 light cycle at 30-70% humidity, ad-libitum water, and standard chow diets (LabDiet, St. Louis, MO).

### Mice infection

Twelve to twenty weeks old male and female K18-hACE2 mice were inoculated intranasally with 10^4^ PFU of SARS-CoV-2 in 50 μl of sterile 1X PBS. The inoculations were performed under 1-3% isoflurane anesthesia. Twenty-six mice (6 for WT, 10 for Omi-S, and 10 for Omicron) were enrolled in a 14-day survival study, and another 42 mice (14 for each of the WT, Omi-S, and Omicron viruses) were used for virological and histological analysis of infected lungs. During the survival study, the animals were monitored for body weight, respiration, general appearance, responsiveness, and neurologic signs. An IACUC-approved clinical scoring system was used to monitor disease progression and define humane endpoints. The score of 1 was given for each of the following situations: body weight, 10-19% loss; respiration, rapid and shallow with increased effort; appearance, ruffled fur and/or hunched posture; responsiveness, low to moderate unresponsiveness; and neurologic signs, tremors. The sum of these individual scores constituted the final clinical score. Animals were considered moribund and humanly euthanized in case of weight loss greater than or equal to 20%, or if they received a clinical score of 4 or greater for two consecutive days. Body weight and clinical score were recorded once per day for the duration of the study. For the purpose of survival curves, animals euthanized on a given day were counted dead the day after. Animals found dead in cage were counted dead on the same day. For euthanization, an overdose of ketamine was administered followed by a secondary method of euthanization.

For quantification of SARS-CoV-2 infectious particles in lungs by the plaque assay, lung tissues were collected in 600 μl of RNAlater stabilization solution (ThermoFisher Scientific; #AM7021) and stored at −80°C until analysis. 20-40 mg of tissue was placed in a tube containing 600 μl of OptiMEM and a 5 mm stainless steel bead (Qiagen; #69989) and homogenized in the Qiagen TissueLyser II by two dissociation cycles (1,800 oscillations/minute for 2 minutes) with a one-minute interval between cycles. The homogenate was centrifuged at 15,000 xg for 10 minutes at room temperature and the supernatant was transferred to a new tube. Ten-fold serial dilutions of this supernatant were used for the plaque assay on ACE2/TMPRSS2/Caco-2 cells, as described above.

For IHC and histologic analysis, the insufflated whole lung tissues were inactivated in 10% neutral buffered formalin at a 20:1 fixative to tissue ratio for a minimum of 72h before removal from BSL3 in accordance with an approved IBC protocol. Tissues were subsequently processed, embedded in paraffin and five-micron sections stained with hematoxylin and eosin (H&E) following standard histological procedures. IHC was performed using a Ventana BenchMark Discovery Ultra autostainer (Roche Diagnostics, USA). An anti-SARS-CoV-2 S antibody (Cell Signaling technologies: clone E5S3V) that showed equivalent immunoreactivity against WT and Omicron spike was used to identify virus-infected cells. Negative and positive controls for IHC included blocks of uninfected and SARS-CoV-2-infected Vero E6 cells, respectively.

### Flow cytometry

For flow cytometry, fixed cells were permeabilized in 1x permeabilization buffer (ThermoFisher Scientific; #00-5523-00) and stained with SARS-CoV-2 nucleocapsid antibody (Rockland; #200-401-A50, 1:1,000), followed by donkey anti-rabbit IgG-AF647 secondary antibody (ThermoFisher Scientific; #A-31573). Gating was based on uninfected stained control cells. The extent of staining was quantified using a BD LSR II flow cytometer (BD Biosciences, CA), and the data were analyzed with FlowJo v10.6.2 (FlowJo, Tree Star Inc).

### Immunofluorescence

Immunofluorescence was performed as described in our previous publication^33^. Briefly, virus-infected cells were fixed in 4% paraformaldehyde and permeabilized in a buffer containing 0.1% Triton X-100 prepared in PBS. Following blocking in a buffer containing 0.1% Triton X-100, 10% goat serum, and 1% BSA, the cells were incubated overnight at 4°C with anti-SARS-CoV Nucleocapsid antibody (1:2,000 dilution). The cells were then stained with Alexa Fluor 568-conjugated goat anti-rabbit secondary antibody (1:1000 dilution) (Invitrogen; #A11008) in the dark at room temperature for 1h and counterstained with DAPI. Images were captured using the ImageXpress Micro Confocal (IXM-C) High-Content Imaging system (Molecular Devices) with a 4x S Fluor objective lens at a resolution of 1.7 micron/pixel in the DAPI (excitation: 400 nm/40 nm, emission: 447 nm/60 nm) and TexasRed (excitation: 570nm/80nm, emission: 624nm/40nm) channels. Both channels were used to establish their respective laser autofocus offsets. The images were analyzed using MetaXpress High Content Image Acquisition and Analysis software (Molecular Devices). First, the images were segmented using the CellScoring module. The objects between 7 and 20 microns in diameter and greater than 1800 gray level units in intensity were identified and classified as nuclei. Positive cells were taken as nuclei having TexasRed signal of 1500 gray level units or above within 10 to 20 microns of each nucleus. The remaining objects were set to negative cells. From these objects, the following readouts were measured and used for downstream analysis: Total number of positive and negative cells, total area of positive cells, and integrated intensity in the TexasRed channel for positive cells. To calculate the 50% neutralizing dilution (ND_50_), we performed a non-linear regression curve fit analysis using Prism 9 software (GraphPad).

## ACKNOWLEDGEMENT

We thank Dr. Yoshiharu Matsuura from Osaka University, Japan, for providing the pCSII-SARS-CoV-2 F8 plasmid; the Department of Public Health, Massachusetts, for providing the clinical specimen containing Omicron virus; and the ICCB-Longwood Screening Facility of Harvard Medical School for assistance with IF image acquisition and analysis. This work was supported by Boston University startup funds (to MS and FD), Peter Paul Career Development Award (to FD), and BMBF SenseCoV2 01KI20172A (to AE) and DFG Fokus COVID-19, EN 423/7-1 (to AE). The authors would also like to acknowledge the NIH S10 Shared Instrumentation Grants, S10-OD026983 and SS10-OD030269 (to NAC), for supporting this work. We thank the Evans Center for Interdisciplinary Biomedical Research at Boston University School of Medicine for their support of the Affinity Research Collaborative on ‘Respiratory Viruses: A Focus on COVID-19’.

## AUTHORS’ STATEMENT ABOUT ACKNOWLEDGEMENT

The acknowledgement section has been updated according to the 2021 NIH guidelines for acknowledging federal funding (https://grants.nih.gov/policy/federal-funding.htm). Based on these guidelines, the authors acknowledge support from two NIH Shared Instrumentation Grants, S10-OD026983 and SS10-OD030269. Funds from these grants were used to purchase shared-use equipment for Boston University core facilities, and this equipment was used in the research described here.

## AUTHOR CONTRIBUTIONS

M.S. conceptualized the study. DYC, AHT, DK, CVC, NK, HLC, FD, and MS performed experiments. GL and MUG established and provided the modified CPER system. NAC performed histopathologic and IHC analysis of mouse lungs. SCB and MB provided scientific input and helped secure funds. AH and AE provided BAC harboring the SARS-CoV-2 genome. JHC provided the Omicron isolate. YK provided plasma samples. MS wrote the manuscript, which was read, edited, and approved by all authors.

## EXTENDED DATA FIGURES

**Extended Data Fig. 1:**
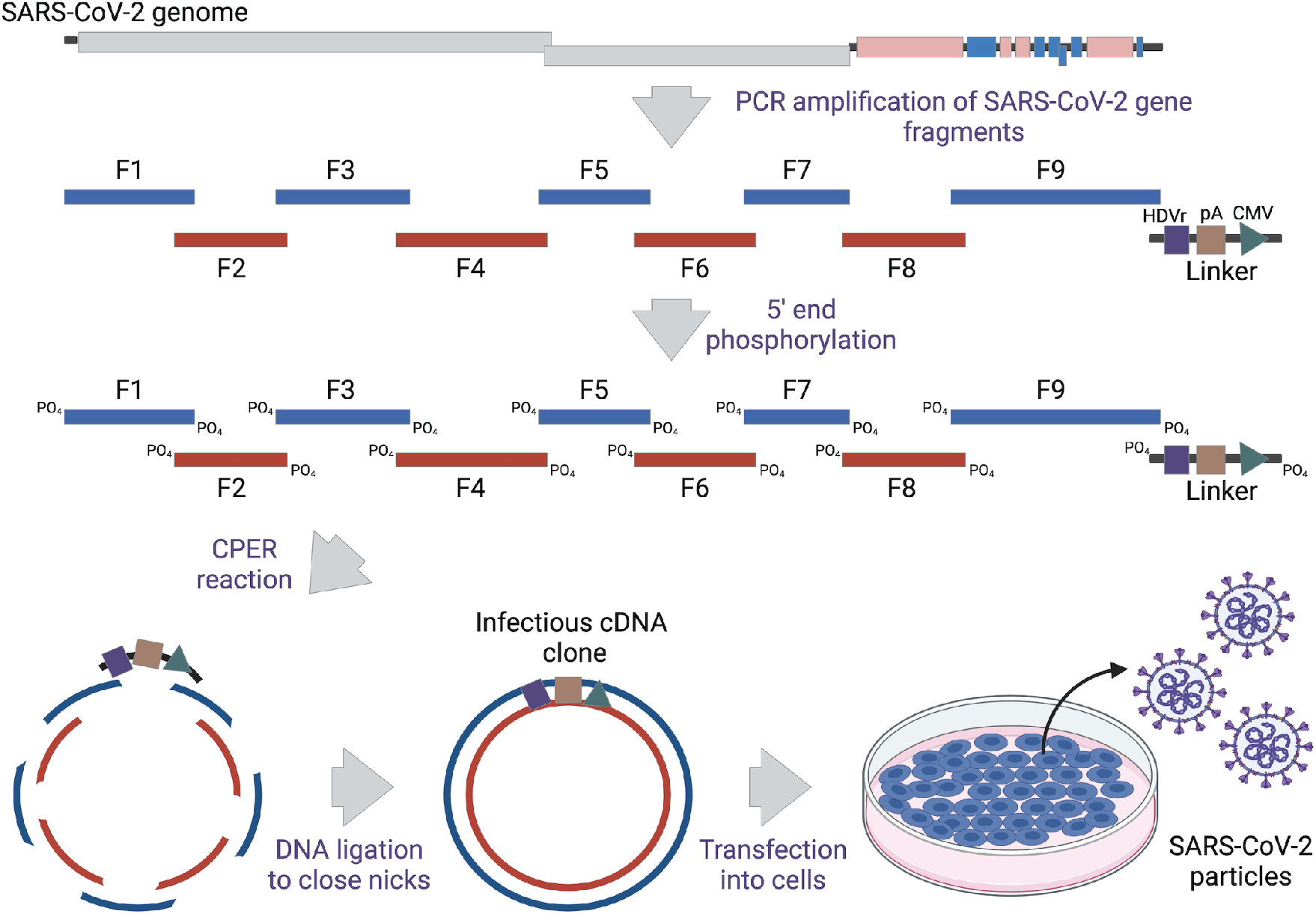
Schematic representation of CPER to generate recombinant SARS-CoV-2. The SARS-CoV-2 genome was amplified into nine overlapping fragments. These fragments and a linker (containing a hepatitis delta virus ribozyme, a poly-A signal, and a CMV promoter) were treated with PNK to phosphorylate 5’ ends. The 5’-end phosphorylated fragments were then stitched together by CPER, and the nicks in the resulting circular DNA molecule were closed by treatment with DNA ligase. The CPER product was transfected into cells to rescue virus particles.

**Extended Data Fig. 2:**
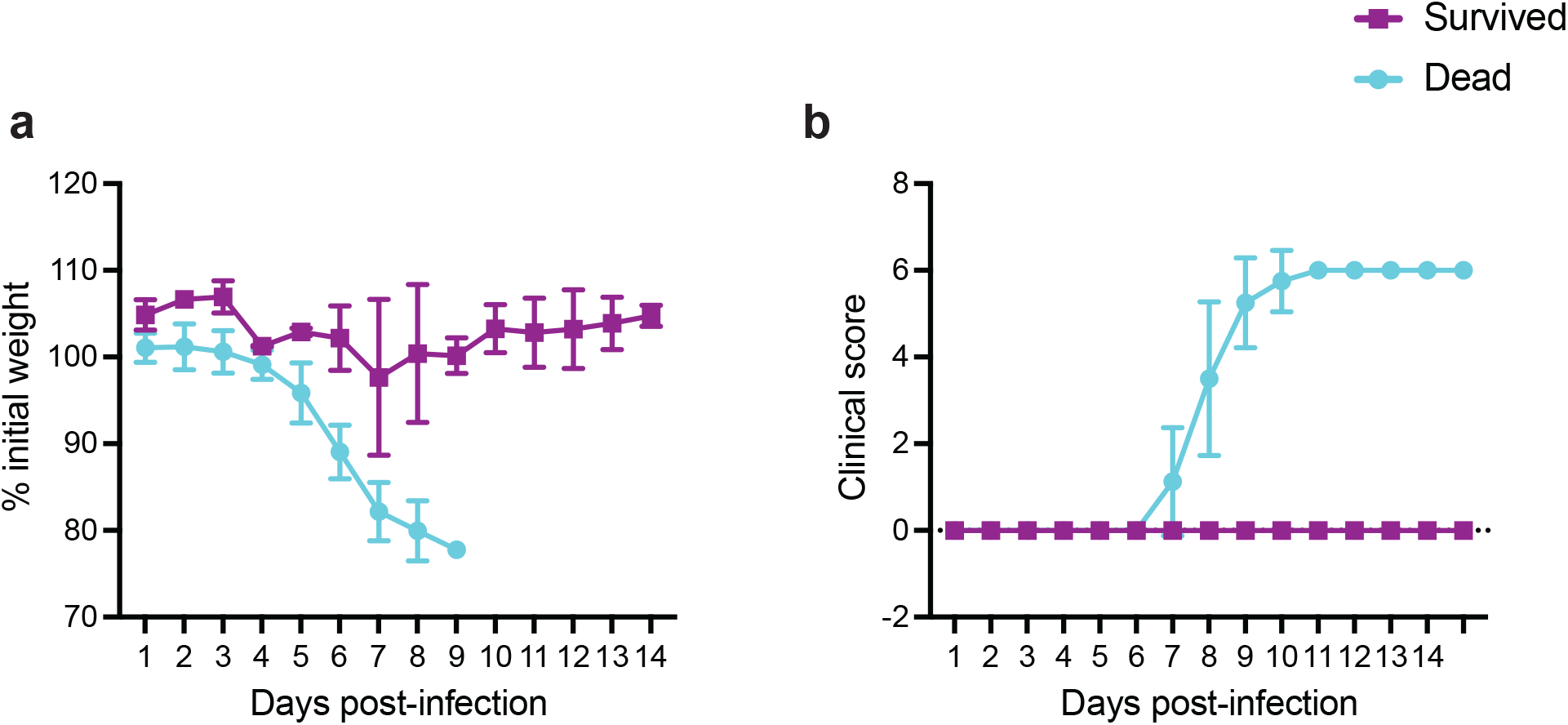
Clinical signs of Omi-S-infected mice. K18-hACE2 mice (*n = 10*) inoculated intranasally with 1 × 10^4^ PFU of Omi-S and described in Fig. 3a-c were monitored for body weight (**a**) and clinical score (**b**). Animals losing 20% of their body weight (8 out of 10) were euthanized. The surviving animals did not show any signs of distress.

**Extended Data Fig. 3:**
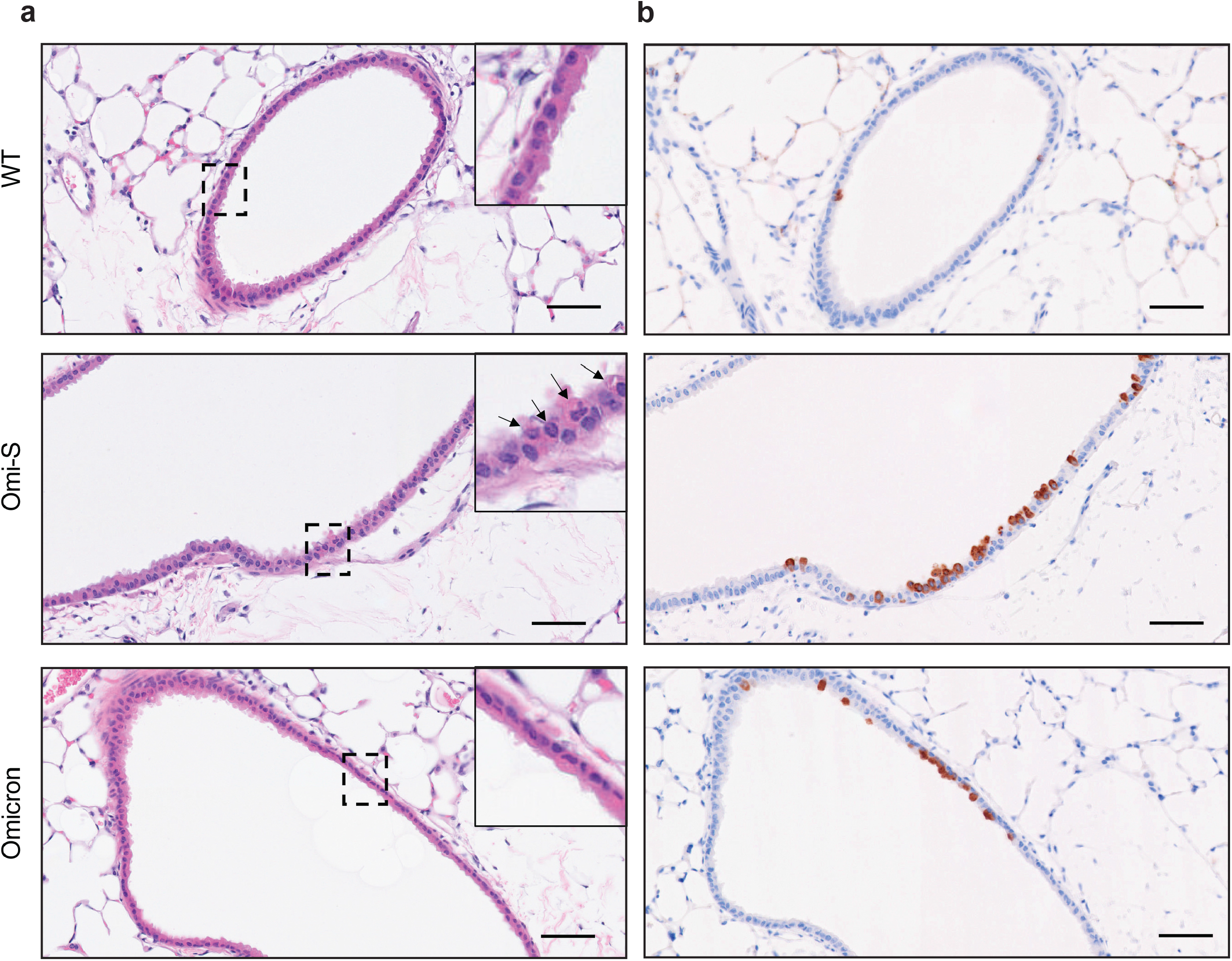
Lung pathology induced by Omi-S. The lungs of the male and female K18-hACE2 mice intranasally inoculated with 1 × 10^4^ PFU of WT, Omi-S, and Omicron were collected at 2 dpi for histological analysis. **a,** Representative images of hematoxylin and eosin (H&E) staining for the detection of bronchiolar damage in the lungs of the infected mice. The bronchiolar epithelial necrosis is shown with arrows. Note that the necrosis was no longer evident at 4 dpi in any cohort. **b,** Immunohistochemistry (IHC) staining for the detection of SARS-CoV-2 S protein in the same area where bronchiolar necrosis was seen. The only bronchiole found to be positive for Omicron is shown. No evidence of necrosis was seen for this bronchiole. (Scale bar = 100 μm).

**Extended Data Fig. 4:**
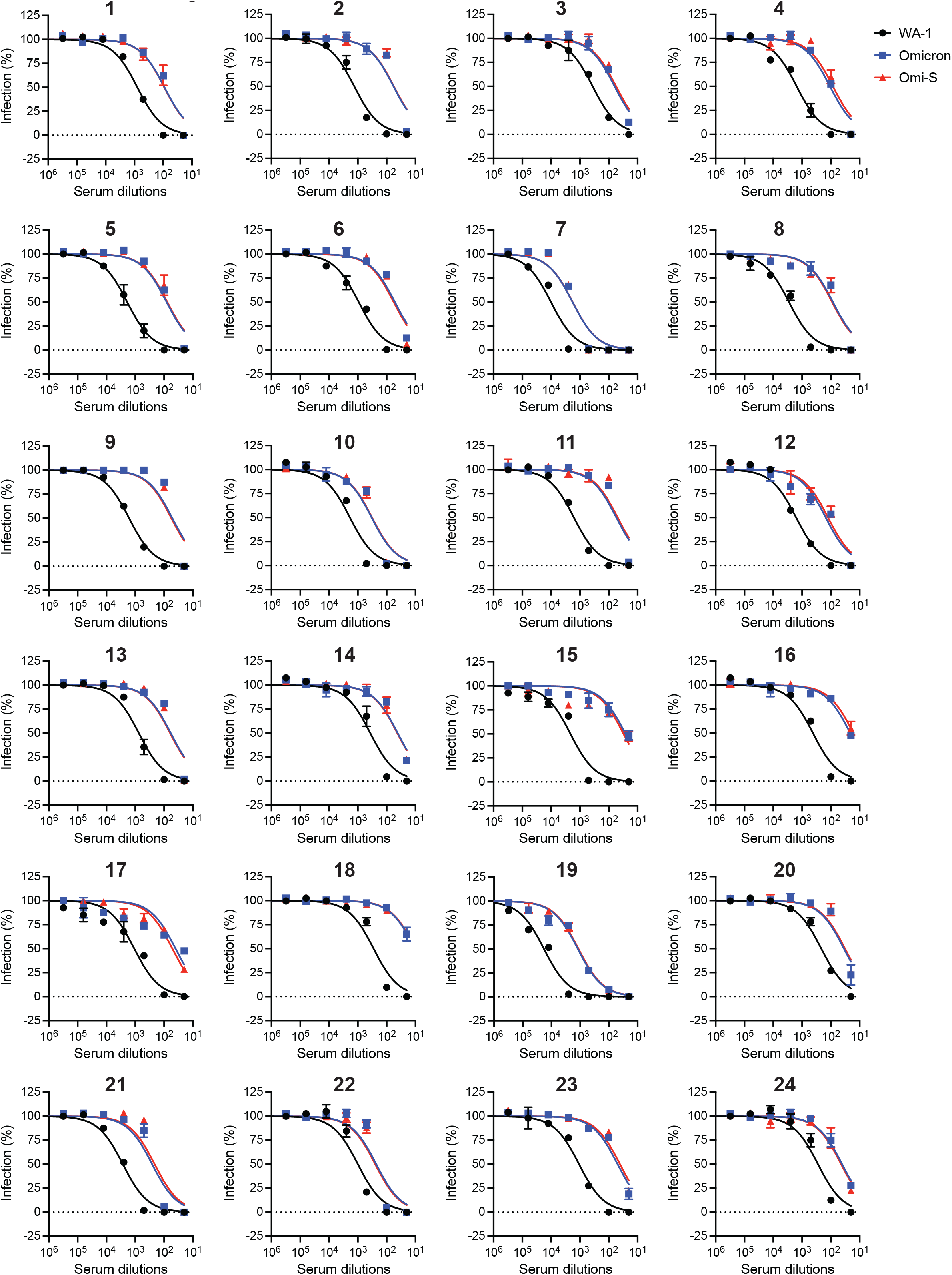
Individual neutralization data. Individual neutralization curves for the data presented in Fig. 4a,b are shown. The data represent the mean ± SD of three technical replicates. The curves were calculated based on a non-linear regression curve fit analysis in Prism. The dotted lines represent the limit of detection.

**Extended Data Table 1:** Overview of serum samples used for the analysis of antibody neutralization of WA1, Omi-S, and Omicron. *Days after the second vaccine shot. **The spike antibody titer was measured by Abbott’s SARS-CoV-2 immunoassays.

| Serum no. | Sex | Race | Age | Days post-vaccination* | Vaccine (Manufacturer) | Spike Ab titer (AU/ml)** |
| --- | --- | --- | --- | --- | --- | --- |
| 1 | Male | White | 59 | 18 | mRNA-1273 (Moderna) | 39823.0 |
| 2 | Male | Black | 26 | 37 | mRNA-1273 (Moderna) | 26978.7 |
| 3 | Male | Asian | 55 | 34 | mRNA-1273 (Moderna) | 24880.7 |
| 4 | Male | White | 39 | 32 | mRNA-1273 (Moderna) | 23816.7 |
| 5 | Male | Asian | 45 | 38 | mRNA-1273 (Moderna) | 21659.5 |
| 6 | Male | White | 30 | 32 | mRNA-1273 (Moderna) | 18986.5 |
| 7 | Female | Asian | 47 | 35 | mRNA-1273 (Moderna) | 100000.0 |
| 8 | Female | White | 62 | 47 | mRNA-1273 (Moderna) | 69680.0 |
| 9 | Female | White | 39 | 14 | mRNA-1273 (Moderna) | 54996.6 |
| 10 | Female | White | 38 | 32 | mRNA-1273 (Moderna) | 46494.7 |
| 11 | Female | White | 34 | 30 | mRNA-1273 (Moderna) | 43784.0 |
| 12 | Female | White | 57 | 42 | mRNA-1273 (Moderna) | 42140.5 |
| 13 | Male | Mixed | 28 | 51 | BNT162b2 (Pfizer-BioNTech) | 17623.8 |
| 14 | Male | White | 30 | 54 | BNT162b2 (Pfizer-BioNTech) | 16154.5 |
| 15 | Male | White | 29 | 54 | BNT162b2 (Pfizer-BioNTech) | 14261.5 |
| 16 | Male | Asian | 48 | 48 | BNT162b2 (Pfizer-BioNTech) | 10593.6 |
| 17 | Male | White | 46 | 60 | BNT162b2 (Pfizer-BioNTech) | 9752.3 |
| 18 | Male | White | 31 | 53 | BNT162b2 (Pfizer-BioNTech) | 8715.2 |
| 19 | Female | White | 55 | 52 | BNT162b2 (Pfizer-BioNTech) | 100000.0 |
| 20 | Female | White | 43 | 47 | BNT162b2 (Pfizer-BioNTech) | 44385.4 |
| 21 | Female | White | 56 | 48 | BNT162b2 (Pfizer-BioNTech) | 39998.5 |
| 22 | Female | Mixed | 44 | 49 | BNT162b2 (Pfizer-BioNTech) | 31141.9 |
| 23 | Female | White | 56 | 50 | BNT162b2 (Pfizer-BioNTech) | 25969.6 |
| 24 | Female | White | 55 | 51 | BNT162b2 (Pfizer-BioNTech) | 23539.1 |

